# Prophages block cell surface receptors to ensure survival of their viral progeny

**DOI:** 10.1101/2024.03.05.583538

**Authors:** Véronique L. Taylor, Megha Shah, Pramalkumar H. Patel, Ahmed Yusuf, Cayla M. Burk, Zemer Gitai, Alan R. Davidson, Matthias D. Koch, Karen L. Maxwell

## Abstract

In microbial communities, viruses compete for host cells to infect, and thus evolved diverse ways to inhibit their competitors. One mechanism is Superinfection exclusion (Sie), whereby a virus that has established an infection prevents a secondary infection. We identified a *Pseudomonas* prophage Sie protein that alters pilus dynamics through the pilus assembly chaperone, PilZ. This protein, known as Zip for Pil<u>Z i</u>nteracting <u>p</u>rotein, does not abrogate pilus activity, but fine tunes it, providing strong phage resistance without a fitness cost. This tuning is modulated through quorum sensing, which coordinates Zip production in concert with bacterial cell density to ensure maximal protection when bacterial populations are at the highest risk of phage infection. Most notably, Zip activity prevents internalization and destruction of phage progeny. We refer to this as the “anti-Kronos effect” after the Greek god who devoured his own children and show that it is a conserved feature of diverse prophage-encoded Sie systems.

## Introduction

The ability of an established viral infection to inhibit subsequent infection by a closely related virus can have profound effects on viral evolution and the structure of viral populations^1^. This phenomenon, known as Superinfection exclusion (Sie), was first characterized in phages, the viruses that infect bacteria^2^. It has subsequently been described for a wide range of viruses that infect animals, plants, and bacteria^3–8^. While genes that endow Sie are a common feature of many viruses, and detailed mechanisms of activity have been determined for many systems, their evolutionary significance has remained largely unclear. Several theories have been put forth – one possible advantage is to protect the infected cell from other viruses that would compete for host cell resources. Alternatively, it has been proposed that Sie may permit cooperation between the same or closely related viruses, with the exclusion of related viruses benefiting the community by allowing them to efficiently search out uninfected cells^9^.

Bacterial viruses can be divided into two major groups, lytic and temperate. Lytic phages infect bacteria and immediately reproduce, destroying the host and releasing the viral progeny. As a result, lytic phages are transmitted horizontally following death of the infected bacterium. By contrast, temperate phages can switch between lytic and lysogenic lifestyles and thus can be transmitted both horizontally and vertically. During lysogeny, temperate phages exist in a quiescent state inside the cell, known as a prophage, and are passively replicated along with the bacterial cell. In this state, the survival of the prophage depends on the survival of the bacterial host cell. As a result, prophages often express genes that increase the fitness of the bacterial host, such as Sie proteins that protect against further phage infection^4,10–12^.

Phages have been shown to mediate superinfection exclusion via mechanisms that inhibit the adsorption of competing phages to the host cell surface or prevent the successful injection of their genomes across the bacterial cell envelope. For example, in *E. coli,* phage T5 encodes a lipoprotein known as Llp that is produced early in infection and prevents further adsorption events by blocking the cell surface receptor protein, FhuA^13^. Phage T4 encodes two proteins, Imm and Sp, that block superinfection by T-even phages by inhibiting phage lysozyme activity, which prevents DNA transfer across the cell envelope^14^. Phage HK97 expresses a small protein that localizes to the inner membrane and inhibits the DNA injection process of superinfecting phages^15,16^. Other phages have been shown to express Sie proteins that block phage DNA transfer across the cell membrane^17–19^ or mask the host cell receptor via serotype conversion^20,21^.

Phages that infect *Pseudomonas aeruginosa* predominantly recognize two cell surface receptors – lipopolysaccharide and the type IV pilus ^23–25^. Sie proteins that confer phage resistance through alterations of each of these receptors have been identified^12,26,27^. For example, *P. aeruginosa* phage D3 encodes three proteins, an α-polymerase inhibitor, a β-polymerase, and an *O*-acetylase, that work together to inhibit formation of O5 B-band lipopolysaccharide, replacing it with β-linked B-band lipopolysaccharide, and modifying the O-antigen via addition of *O*-acetyl groups to mediate superinfection exclusion^25,26^. DMS3-like phages have been shown to encode Sie proteins that interact with the pilus ATPase extension motor PilB, preventing pilus assembly on the cell surface and thereby protecting the prophage-containing cell against superinfection^28,29^. As many *P. aeruginosa* phages rely on the type IV pilus for infection, pilus inhibition provides a useful mechanism for blocking further phage infection. However, this is a double-edged sword; while the disruption of pilus assembly protects against superinfection, *P. aeruginosa* that lack type IV pili are unable to adhere to surfaces and establish beneficial biofilms or escape unfavourable environments^30–32^. How prophages that inhibit pilus assembly to provide superinfection exclusion balance the fitness advantage of phage resistance with the potential cost associated with the loss of the pilus is unknown.

A previous study of *P. aeruginosa* phages identified a protein encoded by phage JBD26 that abrogated twitching motility and endowed strong superinfection exclusion activity against all phages that require the type IV pilus for infection^22^. This gene, *61*, was shown to be highly expressed from the prophage during late exponential growth^27^. However, in contrast to the complete abrogation of twitching motility observed when gene *61* was expressed from a plasmid, expression from its natural context in the JBD26 prophage decreased twitching motility only ∼20% as compared to wild-type PA14, while still providing very strong phage resistance^22,27^. This paradox of high resistance with minimal effects on twitching motility motivated us to investigate this system in detail to determine how the prophage manipulates the bacterial cell to increase fitness via phage resistance while avoiding the evolutionary cost of loss of the pilus. This work also allowed us to address the question of why providing superinfection exclusion at the bacterial cell surface is so advantageous for temperate phages.

In this study, we show that gene *61* provides strong superinfection exclusion activity by modulating pilus assembly through an interaction with the type-IV pilus biosynthesis protein PilZ. We found that expression of gene *61*, which we call *zip* for Pil<u>Z i</u>nhibitory <u>p</u>rotein, is driven by the LasR quorum sensing regulator, providing the first example of a prophage tapping into the host quorum sensing pathway to regulate phage defence. We show that *zip* expression from the prophage not only protects against other phages that use the pilus for infection, but strikingly has a strong protective effect on phage progeny produced by the bacterial community through spontaneous prophage induction. We call this the “anti-Kronos” effect; in the absence of Zip, phage progeny are effectively lost due to reinfection of the lysogen. This cooperative trait endowed by Zip allows phage progeny to efficiently avoid cells already containing a prophage in their search for new hosts to infect. We further show that this trait of prophage resistance at the cell surface to prevent loss of viral progeny is a conserved feature of temperate phages that use diverse cell surface receptors. This work provides important new insight into the evolutionary advantage resulting from lysogens inhibiting self-infection through the activity of Sie proteins.

## Results

### Zip binds to PilZ

In a previous study, we identified a protein expressed from the host genome-integrated form (prophage) of *Pseudomonas aeruginosa* phage JBD26 that decreased twitching motility and mediated strong defence against all phages that require the type IV pilus for infection. This protein, which we call Zip (Pil<u>Z i</u>nteracting <u>p</u>rotein; YP_010299255), was shown to be highly expressed from the prophage during late exponential growth^27^. To determine the mechanism by which Zip provides phage defence, we first sought to identify its bacterial target. As Zip expression inhibited twitching and mediated strong resistance to phages that require the type IV pilus for infection, we performed a bacterial two-hybrid assay^33^ with a library expressing components of the *P. aeruginosa* type IV pilus. This library included major and minor pilins, chaperones, scaffold, and motor protein genes. We detected a strong interaction between Zip and PilZ (PA2960; Figure S1A). Previously, a *pilZ* deletion was shown to prevent type IV pilus assembly, thereby abrogating twitching motility and increasing phage resistance^34^. These phenotypes are consistent with those observed with Zip expression, suggesting that PilZ is a biologically relevant target.

*P. aeruginosa* type IV pili can be classified into five different families based on the sequence of the major pilin subunit, PilA, and the presence or absence of associated major and minor pilin proteins^35^. PilZ shares 100% sequence identity across these five families, suggesting that Zip activity should inhibit all of them. To test this, we expressed Zip from a plasmid in five strains that each encode a different pilus type and determined that it abrogated twitching motility across all five families (Figure 1A). We next overexpressed PilZ from a plasmid in cells containing a JBD26 prophage to see if this could rescue the ability of phages to plate by ensuring that excess PilZ was present in the cell to overcome the Zip-mediated inhibition. We found that overexpression of PilZ in the lysogen increased phage susceptibility by >10^4^-fold (Figure 1B), providing further evidence that the highly conserved PilZ is the target of Zip activity.

**Figure 1.**
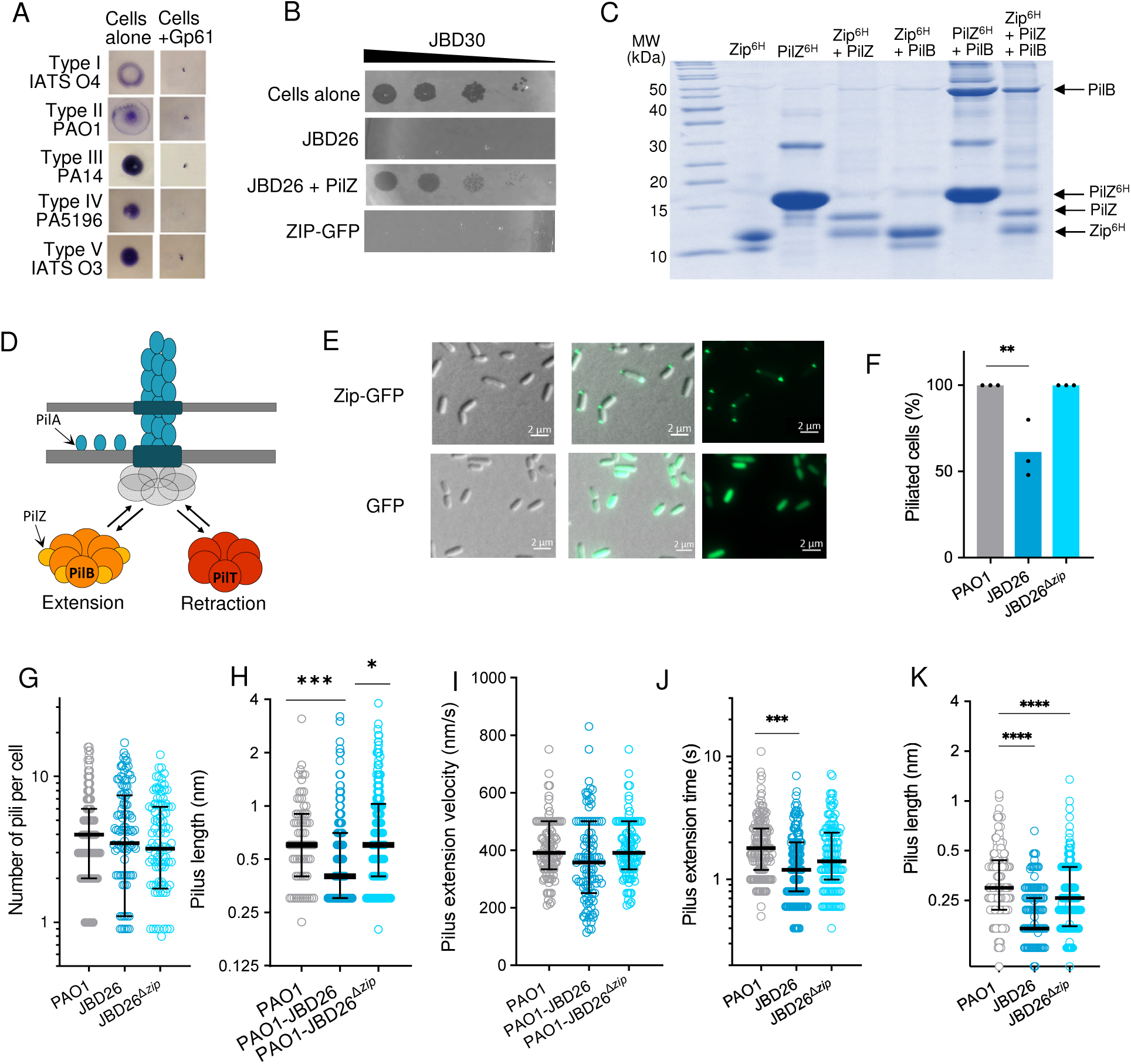
Zip targets a conserved pilus chaperone, PilZ. **(A)** Twitching motility of cells expressing Zip from a plasmid. **(B)** Ten-fold serial dilutions of phage JBD30 plated on laws of PAO1, the JBD26 lysogen, JBD26 lysogen expressing PilZ from a plasmid, and PAO1 expressing Zip-GFP. A representative image of three replicates is shown. **(C)** SDS-PAGE gel analysis of co-purifications of Zip, PilZ and PilB stained with Coomassie blue. **(D)** Pilus assembly and retraction results from the activity of the PilB assembly ATPase, which incorporates PilA monomers into the growing filament, and the PilT ATPase, which disassembles the filament and causes pilus retraction. **(E)** Fluorescence microscopy of cells expressing GFP-tagged Zip (top), or GFP alone (bottom). **(F)** Percent of cells that produce at least one pilus in 30 secs after overnight growth. Data are representative of three biological replicates. Significance of a one-way ANOVA (** p < 0.005) shown. (**G**) – (**J**) Pilus dynamics of mid-log cells (OD = 0.3). (**G**) Total number of pili individual cells make in a 30 second time interval. (**H**) Maximum length of individual pili. (**I**) The extension velocity of individual pili. (**J**) Total time individual pili spend extending. Significance of a two-tailed Mann-Whitney test (*** p < 0.05) (**K**) Length of pili individual cells make in a 30 second time interval at high cell density (overnight culture). Results of a one-way ANOVA (**** p < 0.0001).

PilZ, a known regulator of type IV pilus function, has previously been shown to bind the PilB assembly ATPase protein that is required for type IV pilus biogenesis. This interaction is thought to stabilize the PilB N-terminal domain and allow a hexamer of PilB to efficiently localize to the pilus scaffold complex^36,37^. Once attached to the pilus scaffold, the PilB hexamer hydrolyses ATP and provides the energy to incorporate hundreds of PilA subunits into the extending pilus. To determine if Zip affects the ability of PilZ to chaperone PilB, we used protein co-purification experiments to examine the complexes. First, we confirmed the direct interaction between Zip and PilZ by co-expressing His-tagged Zip with untagged PilZ in *E. coli* and performing a Ni-affinity chromatography co-purification experiment. We found that untagged PilZ co-eluted with the His-tagged Zip, confirming direct protein binding (Figure 1C). We next co-expressed His-tagged PilZ and untagged PilB and detected a stable complex (Figure 1C). To determine if Zip interferes with this interaction, we co-expressed His-tagged Zip together with untagged PilZ and PilB and performed Ni-affinity purification. We found that all three proteins co-purified, suggesting the formation of a tripartite complex (Figure 1C). To further characterize this complex and determine if Zip binds directly to PilB or if this interaction is mediated through PilZ, we co-expressed His-tagged Zip with untagged PilB and found that they did not co-purify (Figure 1C). Together, these data imply that Zip interacts with PilZ, and that Zip does not prevent PilZ from binding to PilB.

### Zip localizes to the cell poles and makes shorter pili

The PilZ-PilB interaction is known to allow efficient localization of the PilB ATPase to the pilus scaffold complex located at the poles of the cell^37^. The activity of Zip could interfere with this localization, or it could affect the ability of the PilB motor to incorporate the PilA subunits into the extending pilus filament (Figure 1D). To determine if Zip localizes to the poles of the cell where the pilus complex is known to assemble, we expressed green fluorescent protein (GFP)-tagged Zip with its native promoter from a plasmid in PAO1 and examined its localization using fluorescence microscopy. First, to ensure that the fusion of GFP to the C-terminus of Zip did not interfere with its activity, we assessed twitching motility and phage resistance of cells expressing the fusion protein. We found that Zip-GFP expression in PAO1 resulted in a decrease in twitching of approximately 20% as compared to wild-type PAO1, a level comparable to the twitching inhibition observed with the JBD26 lysogen (Figure S1B). Zip-GFP expression also provided strong phage resistance (Figure 1B) implying that the GFP tag did not inhibit activity. Examination of these cells using fluorescence microscopy revealed that Zip-GFP accumulated at the poles of the cells while GFP alone was diffusely located throughout the cytoplasm (Figure 1E). These data show that Zip localizes to the poles of the cells where the type IV pilus machinery is known to assemble.

While the co-purification experiments confirmed that PilZ is the target of Zip binding, it was not clear how this interaction would affect type IV pilus assembly. To gain insight into Zip activity in the context of the prophage, we used fluorescence microscopy to directly examine pili on the surface of cells that contain either a wild-type JBD26 prophage or one lacking *zip* (JBD26^Δ^*^zip^*). In these experiments, the major pilin subunit (PilA) was fluorescently labelled with thiol-reactive maleimide dye Alexa88-mal^38^. We first compared the number of pili present on the surface of the cells at mid-log phase (OD ∼0.3) and following overnight growth. We found no significant difference between wild-type PAO1, the JBD26 lysogen or the JBD26^Δ^*^zip^* lysogen during early exponential phase; all cells in each of the three samples were piliated, with an average of ∼4 pili assembled per cell per 30 second increment (Figure 1G). By contrast, following overnight growth we found that ∼60% of JBD26 lysogens had no pili on their surface, while 100% of the wild type PAO1 and the JBD26^Δ^*^zip^* lysogen were piliated (Figure 1F). Notably, there were also fewer pili on the surface of these cells than at mid-log phase growth, with more than half of the PAO1 and JBD26^Δ^*^zip^* lysogen cells displaying only a single pilus as compared to four at the earlier timepoint.

To gain additional insight into how Zip activity affected pilus assembly, we measured the length of pili on the surface of cells, as well as their extension and retraction velocities. Using the early exponential phase cells where all cells displayed at least one pilus, we first calculated the distribution of pilus lengths. We found that the JBD26 lysogen had a median length of 0.4 μm, while wild-type PAO1 and JBD26^Δ^*^zip^* lysogen showed median lengths of 0.6 μm (Figure 1H). To determine why the pili were shorter on average in the JBD26 lysogen, we next examined extension dynamics of individual pili. While the extension velocity was similar across the three strains (Figure 1I), the extension time was shorter for the JBD26 lysogen (Figure 1J). The current model for pilus assembly suggests that only one type of motor, PilB extension or PilT retraction, can be bound to the pilus machinery at a time^39^. In addition, the binding and unbinding of these motors is stochastic, with the duration of each pilus extension event proportional to how quickly the PilB extension motor becomes unbound^40^. Thus, our data suggest that Zip activity destabilizes the interaction of PilB with the pilus machinery, causing the extension motor to disassociate from the scaffold earlier and thereby generate shorter pili. As we had evidence that Zip activity was increased following overnight growth, with only 60% of JBD26 lysogens displaying pili in the 30 second timeframe we examined, we also examined the length of the pili on these cells (Figure 1K). We found that the JBD26 lysogen pili were shorter on average, at a length of 0.2 μm as compared to 0.4 μm for PAO1. Together, these data imply that Zip activity does not prevent pilus assembly, but instead modulates it in a way that decreases the length of pili produced. In addition, at high cell density Zip expression causes complete loss of pili in almost half the bacterial community (Figure 1F).

### Zip expression is controlled by LasR, the principal quorum sensing regulator

Our previous work showed that *zip* is the most highly expressed gene in the JBD26 prophage during late exponential phase^27^. As it appeared to be expressed independently from other prophage genes, we looked for a potential promoter and discovered a 74-nucleotide non-coding region upstream of the gene. To determine if this sequence includes a promotor that drives *zip* expression, we cloned it into the pQF50 reporter plasmid^41^, which placed it upstream of *lacZ* and allowed its activity to be monitored via β-galactosidase activity. We transformed this plasmid into wild-type PA14 and observed robust β-galactosidase activity at 18 hours (Figure 2A). As this experiment was done in the absence of a JBD26 prophage, it suggested that expression from the JBD26 *zip* promoter not reliant on a phage transcriptional regulator. To determine if a bacterial transcriptional regulator was mediating expression from the *zip* promotor, we assayed β-galactosidase activity in PA14 strains containing single gene deletions of various transcriptional regulators. We found that a culture of the *lasR* mutant (Δ*lasR*), the principal regulator of quorum sensing, showed greatly decreased β-galactosidase activity following overnight growth (Figure 2A). Expression in the *rhlR* mutant was partially attenuated (Figure 2A). As RhlR and LasR are known to activate transcription by binding to a DNA sequence called the las-box with different affinities, finding that both regulators affected *zip* transcription but to different degrees was consistent with previous studies^42–44^. By contrast, mutants in other quorum regulators such as *vfr* and *pilR* did not show a significant difference in β-galactosidase activity as compared to wild-type PA14, suggesting that they are not involved in activation of gene expression associated with this promoter sequence. To provide additional evidence that LasR was driving β-galactosidase expression, we transformed the reporter construct into a QteE deletion strain (Δ*qteE*). QteE reduces the stability of LasR, and its absence allows more LasR to accumulate at lower cell density^45^. We observed increased β-galactosidase activity during exponential growth phase in the Δ*qteE* mutant as compared to wild-type PA14 (Figure 2B), consistent with LasR activating expression from the promoter. These data imply that the *zip* promoter is controlled by the bacterial quorum sensing regulator, LasR. To confirm that LasR controls *zip* expression from the prophage, we tested the ability of the JBD26 prophage to resist infection by the pilus-dependent phage, JBD30; while the JBD26 prophage in wild-type PA14 robustly resisted JBD30 infection, it showed >1,000-fold increased susceptibility to JBD30 infection in the Δ*lasR* strain (Figure 2C). These data show that *P. aeruginosa* prophages exploit the host cell quorum sensing pathway to drive expression of anti-phage defence.

**Figure 2.**
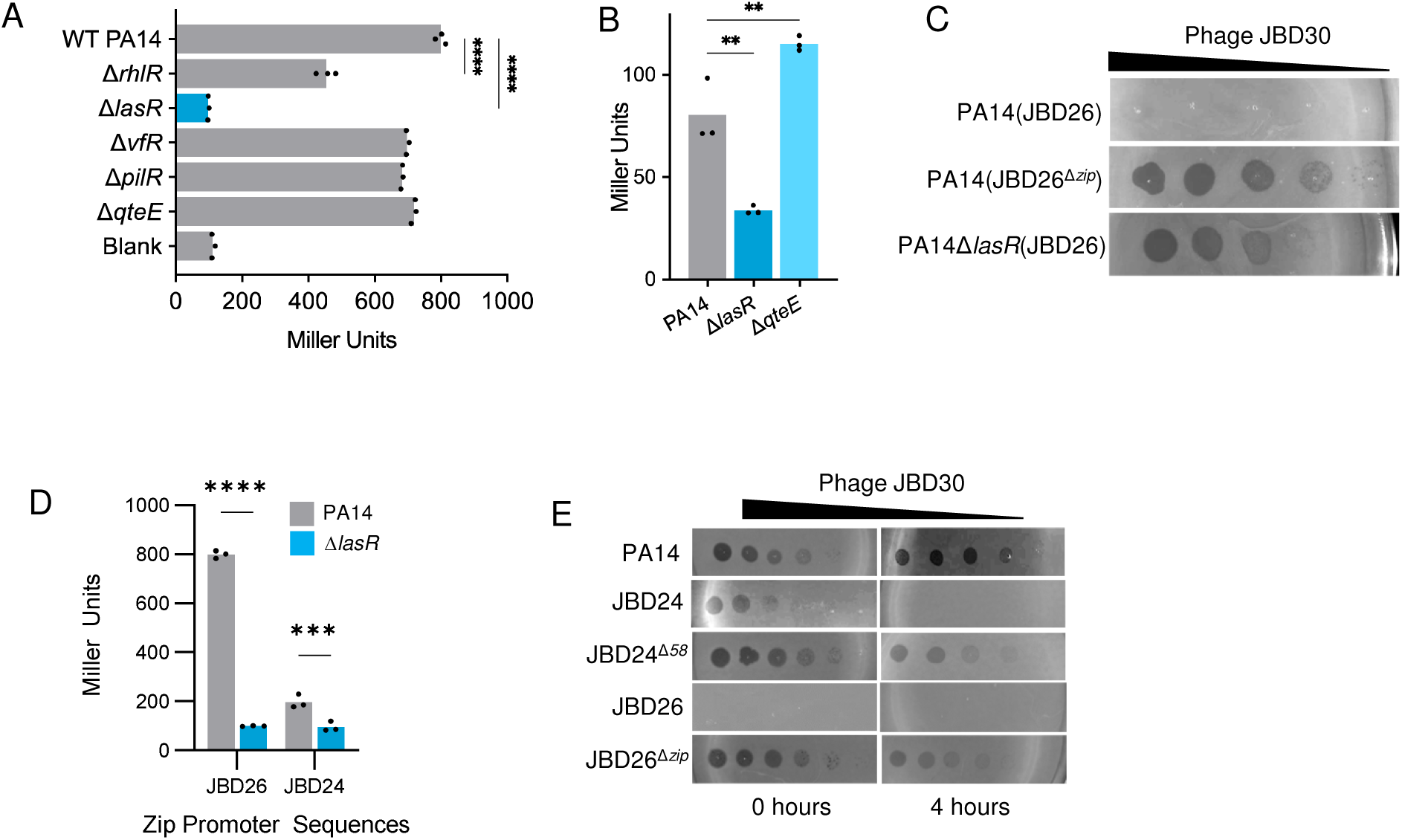
LasR drives Zip expression. **(A)** β-galactosidase activity resulting from gene expression driven by the Zip promoter. Single gene deletions in various PA14 transcriptional regulators are shown. Three independent biological replicates are shown. Statistical significance was calculated using a one-way ANOVA (**** p < 0.0001). **(B)** β-galactosidase activity of the Zip promoter measured during late-exponential growth in PA14, ΔlasR and ΔqteE strains. Three biological replicates are shown. Significance by a one-way ANOVA (** p < 0.005). **(C)** Serial dilutions of phage JBD30 spotted on a wild type JBD26 lysogen, a JBD26^Δ*zip*^ lysogen, and a wild type lysogen in a Δ*lasR* background. A representative image of three biological replicates is shown. **(D)** β-galactosidase activity driven by the *zip* promoter from JBD26 and JBD24 in PA14 and the Δ*lasR* strain. Three biological replicates are shown. Significance by a two-way ANOVA (*** < 0.005, **** <0.0001). **(E)** Serial dilutions of phage JBD30 spotted on lawns either immediately after pouring (0 hours) or following incubation to allow cell growth (4 hours). An image representative of three independent biological replicates is shown.

We had previously noted that a JBD24 lysogen, which is closely related to JBD26 and encodes a homologue of Zip that shares 95% sequence identity (Gp58; YP_007392821), was sensitive to infection by most pilus-dependent phages^22^. A sequence alignment revealed that the promoter region upstream of JBD24 gene *58* differed from the JBD26 sequence upstream of *zip*, sharing 71% sequence identity (Figure S2A). To determine if this promoter sequence was also responsive to LasR we cloned it into the pQF50 reporter plasmid upstream of *lacZ* and assayed its activity using a β-galactosidase assay. This promoter showed much weaker activity than the *zip* promoter from JBD26, but still appeared responsive to LasR (Figure 2D). This weaker promoter activity provides an explanation for the lower superinfection exclusion activity previously noted for the JBD24 lysogen.

To determine if quorum sensing might upregulate expression of Zip and provide JBD24 with greater protection from superinfection at high cell density, we applied a phage JBD30 lysate onto a JBD24 lysogen bacterial lawn that was pre-incubated at 30 °C for several hours to allow the quorum sensing system to activate. We found that JBD30 was able to form plaques on the JBD24 lysogen when phages were applied immediately. By contrast, when the JBD24 lysogen lawn was grown for 4 hours before the phages were applied, complete inhibition of plaquing was observed (Figure 2E), suggesting that defence was upregulated at high cell density. To determine if this effect was due to the activity of the Zip protein, we created a mutant phage lacking gene *58* (JBD24^Δ*zip*^) and repeated the experiment. We found that the ability of the JBD24^Δ*zip*^ lysogen to resist infection by phage JBD30 was greatly diminished when the phage lysate was applied both immediately and after 4 hours of cell growth (Figure 2E). These data show that the JBD24 Zip protein does provide effective Sie protection, and that this activity is activated at high cell density. This contrasts with JBD26, where Zip defence is active at both low and high cell densities (Figure 2E).

### The Kronos effect: lysogens destroy their viral progeny in the absence of Zip

JBD26 prophages spontaneously induce such that an overnight culture of a JBD26 lysogen grown under normal conditions contains abundant viral particles to a level of ∼10^6^ plaque forming units (pfu)/mL. Remarkably, we discovered that an overnight culture of a JBD26^Δ*zip*^ lysogen contained fewer than 500 pfu/mL. To investigate this phenomenon, we monitored the number of phage progeny produced by lysogens of JBD26 and JBD26^Δ*zip*^ over time. To do this, we first removed any free phages from overnight cultures of JBD26 and JBD26^Δ*zip*^ by collecting the cells by centrifugation and washing several times. We then grew the cultures and took samples at 3, 6, and 20 hours after the start of cell growth and titered the number of phages at each time point. We found that while JBD26 and JBD26^Δ*zip*^ lysogen cultures accumulated similar numbers of phages in the supernatant at both 3 and 6 hours, after 20 hours the titer of JBD26^Δ*zip*^ was only ∼500 pfu/mL as compared to ∼10^6^ pfu/mL in the wild-type JBD26 lysogen culture (Figure 3A). Notably, between 6 and 20 hours the JBD26^Δ*zip*^ lysogen titer fell from ∼60,000 pfu/mL to 500 pfu/mL, showing that phages were being actively destroyed. To confirm that the observed difference in phage numbers was due to the loss of Zip activity and not a phage induction or replication defect, we treated the two lysogens with mitomycin C, a DNA damaging agent that causes induction of most of the prophages and subsequent cell lysis. We found that both JBD26 and JBD26^Δ*zip*^ prophages produced titers of 10^6^ pfu/mL (Figure 3B), showing that phage replication was not affected by *zip* deletion. The difference in the number of phage progeny at 20 hours was also not due to differences in bacterial population sizes as both JBD26 and JBD26^Δ*zip*^ cultures grew to 4×10^9^ colony forming units (cfu)/mL at 20 hours (Figure 3C). Since both JBD26 and JBD26^Δ*zip*^ lysogen cultures accumulated equal numbers of phages at 6 hours, we postulated that the loss of phages at the 20-hour timepoint might be due to the JBD26^Δ*zip*^ lysogens not being able to block infection at the cell surface via the activity of Zip, as does the wild-type JBD26 lysogen. This could lead to reinfection by the phage progeny spontaneously produced by the bacterial lysogen. To determine if this were true, we repeated the growth experiment with the same lysogens containing a plasmid expressing Zip. We found that when Zip was supplied from a plasmid, the JBD26 and JBD26^Δ*zip*^ lysogens displayed equal numbers of free phages in overnight cultures. This shows that the loss of Zip activity is the reason for the low titer observed in the JBD26^Δ*zip*^ lysogen (Figure 3D).

**Figure 3.**
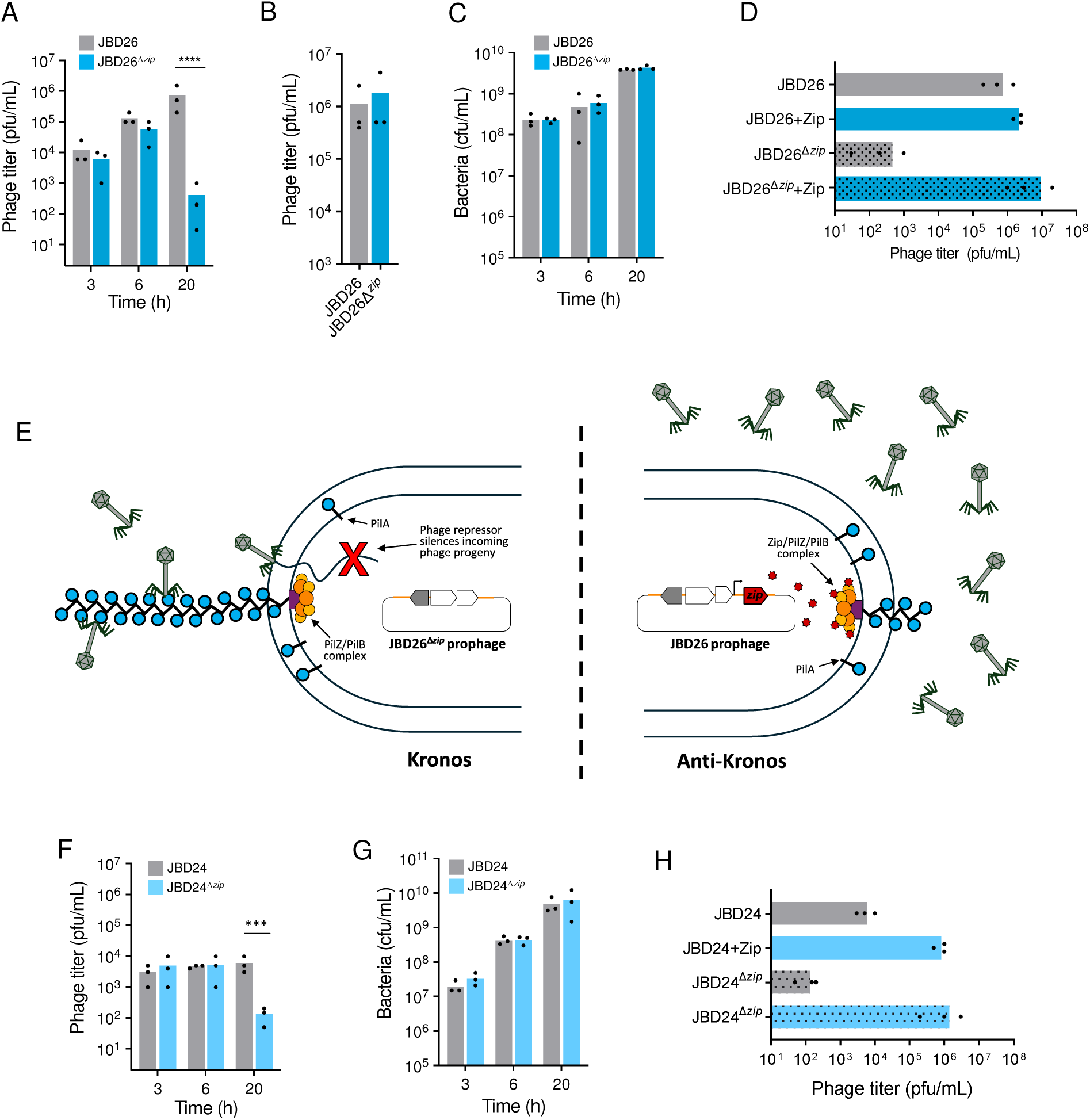
The anti-Kronos effect promotes survival of viral progeny. Phage titers **(A)** spontaneously produced by JBD26 and JBD26 ^Δ^ *^zip^* lysogens over time and **(B)** following mitomycin C treatment. **(C)** Number of bacterial cells present in JBD26 and JBD26^Δ*zip*^ lysogen cultures over time. **(D)** Phage production by JBD26 and JBD26^Δ*zip*^ lysogens in the absence and presence of Zip supplementation from a plasmid. **(E)** The anti-Kronos effect provides the JBD26 lysogen with superinfection exclusion activity that vastly increases the number of phage progeny that accumulate in the community. **(F)** Phage titers spontaneously produced by JBD24 and JBD24 ^Δ^ *^zip^* over time and **(G)** following mitomycin C treatment. **(H)** Phage production by JBD26 and JBD26^Δ*zip*^ lysogens in the absence and presence of Zip supplementation from a plasmid. For all panels, data are representative of three independent biological replicates. Statistical significance was measured using one-way ANOVA, significant p-values are noted (****p < 0.0001, ***p<0.005).

Taken together, these data indicate that Zip performs a crucial function within a lysogenic culture by preventing phage particles produced through spontaneous prophage induction from adsorbing to and injecting their DNA into other lysogenic cells (Figure 3E). As only a small percent of cells in the community are spontaneously producing phage, the chance of the progeny adsorbing to and infecting other lysogens is high. Even though these phage infection events would not kill the lysogens due to immunity conferred by the prophage-expressed repressor protein, which represses phage transcription when the genome enters the cell, the unproductive infections lead to abundant loss of viable phage particles. This effect is shown by the low number of phage progeny observed in the JBD26^Δ*zip*^ lysogen culture at 20 hours (Figure 3A). Zip expression prevents these unproductive infections by preventing phage adsorption to the pilus, and thereby avoiding futile injection of phage DNA into lysogenic cells (Figure 3E). This phenomenon vastly increases the number of free phages in the community that could go on to infect other non-lysogenic hosts. We have named the phenomenon of lysogenic cells destroying their own viral particles through surface adsorption and DNA injection as the “Kronos effect” after the Greek Titan Kronos, who ate his own children. Thus, prophage-encoded proteins that block phage adsorption at the cell surface can be referred to as anti-Kronos factors.

While the JBD24 Zip protein did not endow phage resistance at low cell density (Figure 2E), we wondered if it would show a similar anti-Kronos effect. To investigate this, we assessed the number of free phages present in overnight cultures of JBD24 and JBD24^Δ^*^zip^* lysogens. We found that the wild-type JBD24 lysogen accumulated ∼10,000 pfu/mL, while the JBD24^Δ^*^zip^* lysogen produced only ∼100 pfu/mL (Figure 3F), despite both cultures growing to equal cell densities (Figure 3G). Like JBD26^Δ^*^zip^*, the JBD24^Δ^*^zip^* lysogen showed phage production equal to the wild-type lysogen at 6 hours, and then the phage titer decreased overnight. To determine if higher Zip expression levels would result in the accumulation of more phages in these cultures, we repeated the experiment with the lysogens containing a plasmid expressing Zip. We found that the phage titers of both lysogens increased to ∼10^6^ pfu/mL (Figure 3H), implying that the low titers observed were a result of the Kronos effect. It is interesting to note that the phage titers of JBD24 produced in the presence of plasmid-expressed Zip were equal to the phage titer produced by the wild-type JBD26 lysogen (Figures 3A,F). The lower phage titer naturally produced by the JBD24 lysogen is likely the result of the lower expression levels provided by the native *zip* promoter found in this phage (Figure 2D), resulting in a stronger Kronos effect. These data confirm that the Zip homologue in JBD24 is also required for stable accumulation of free phages in the community.

The *zip* homologues were found in the same genomic position in the closely related phages JBD26 and JBD24. To determine how widespread homologues of this protein are, we performed a PSI-BLAST search starting with the JBD26 Zip sequence. We identified ∼1000 homologues in a variety of *P. aeruginosa* strains and phages, with 73 of the 240 complete *Pseudomonas* phage genomes present in the NCBI database encoding one, suggesting that this is a conserved function. Alignment of these protein sequences showed high conservation among the homologues, with pairwise sequence identities above 80% (Figure S2B). While most phage homologues were found in the same genomic position as *zip*, immediately following the late gene operon, some were found at the other end of the late gene operon, upstream of the small terminase gene (Figure S2C). Regardless of the genomic position, the phage homologues maintained the 5’ upstream region corresponding to the *zip* promoter. These data show that Zip is a conserved feature of *P. aeruginosa* phages and that it provides a widespread mechanism through which lysogens can ensure that phage progeny are protected from loss via the anti-Kronos effect.

### The anti-Kronos effect is conserved and mediated through diverse mechanisms

As the anti-Kronos effect provides such a striking potential evolutionary advantage by greatly increasing the number of phage progeny that can accumulate within a lysogenic community and go on to infect new hosts, we wondered if this effect is widespread and can be mediated by diverse mechanisms. We first examined another pilus-mediated Sie protein, found in *P. aeruginosa* phage LESϕ3. This protein, encoded by gene *50* and referred to here as gp50, interacts with the type IV pilus machinery to block infection by LESϕ3 and other related phages. Gp50 is not related to Zip and inhibits phage attachment by a mechanism that is completely distinct from the Zip mechanism (A. Davidson, manuscript will be uploaded to BioRxiv within the next 2 weeks). Like *zip*, gene *50* has a predicted untranslated region at its 5’ end, suggesting that its expression is likely driven by its own promoter (Figure 4A). To determine if gp50 also protects phage progeny from the Kronos effect, we assessed the number of phages that accumulated in an overnight culture of a mutant LESϕ3 lysogen that lacks gene *50* (LESϕ3^Δ50^). We found that the LESϕ3^Δ50^ lysogen produced an equal number of phage progeny as wild-type LESϕ3 when induced with mitomycin C (Figure 4B, right), showing that phage replication and assembly were not affected by this gene deletion. We next examined the number of phage progeny spontaneously produced by the lysogens at 3, 6, and 20 hours and found that there was a 10^4^-fold decrease in free phage progeny in the LESϕ3^Δ50^ culture at 20 hours (Figure 4B, left). These results show that the anti-Kronos effect is also manifested in this system.

**Figure 4.**
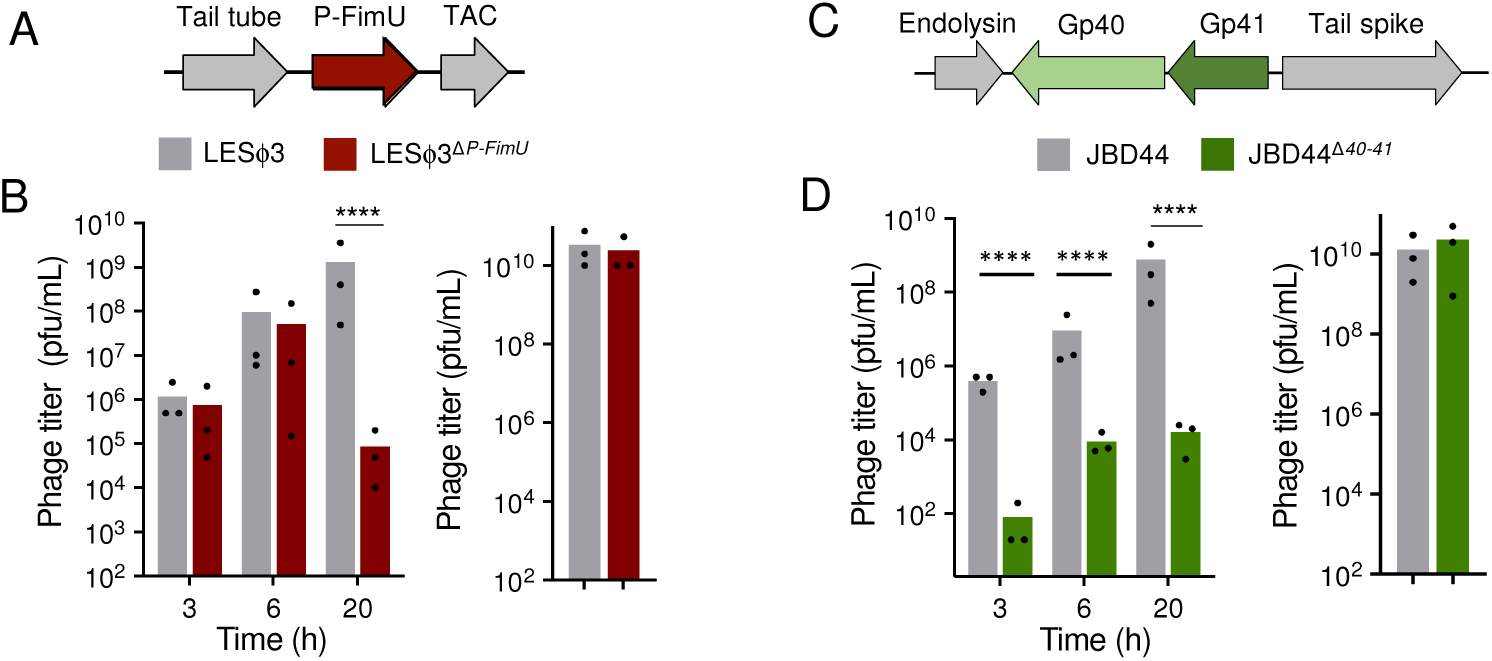
Anti-Kronos systems are widespread. **(A)** Phage LESφ3 encodes a Sie protein between the tail tube and tail assembly chaperone (TAC). **(B)** Phage titers spontaneously produced by LESφ3 and LESφ3^ΔP-FimU^ over time (left) and following mitomycin C induction (right). **(C)** Phage JBD44 encodes a two gene Sie system between the endolysin and tail spike proteins. **(D)** Phage titers spontaneously produced by JBD44 and JBD44^Δ40–41^ over time (left) and following mitomycin C induction (right).

Not all *P. aeruginosa* phages use the type-IV pilus for infection. We wondered if phages that rely on the other major cell surface receptor for phages, lipopolysaccharide (LPS), also modify their surface to protect against the Kronos effect. Phage JBD44, which is known to use LPS as a cell surface receptor^22^, encodes two genes whose activity acetylates the bacterial O-antigen and provides an LPS modification known as serotype O10 to the host in which the prophage resides ^46^. These genes are located at the end of the tail operon, between the tail spike and endolysin genes (Figure 4C). We deleted the genes corresponding to these LPS-modifying enzymes from JBD44 (JBD44^Δ40–41^) and monitored phage production over time. We found that the JBD44^Δ40–41^ lysogen culture accumulated ∼10^4^-fold fewer phages/mL in the culture medium than wild-type JBD44 after overnight incubation (Fig. 4D, left). This was not due to problems with phage replication or assembly of the virion as the mutant phage produced the same number of phage progeny following mitomycin C induction (Figure 4D, right). In contrast to JBD26 and LESϕ3, the Kronos effect is strongly observed at all timepoints. This is likely a result of the different receptors being used. Both JBD26 and LESϕ3 use the type IV pilus, which is not highly expressed in cells growing in liquid medium, and thus does not provide many receptors for phages to bind. By contrast, JBD44 uses LPS, which coats the entire cell and provides an abundance of receptor with which the phages can interact and thereby be lost through the Kronos effect. These data show that JBD44 also expresses anti-Kronos proteins that protect spontaneously produced phage progeny in the community from being lost due to reinfection of the lysogen.

## Discussion

It has long been known that prophages express a variety of proteins that provide them with defence against further phage infection^47–49^. Many of these defences have been shown to act at the cell envelope, blocking the cell surface binding or genome injection of superinfecting phages. Here, we characterize a superinfection exclusion protein that binds the PilZ assembly chaperone and modulates *P. aeruginosa* type IV pilus biosynthesis. The expression of Zip from the JBD26 prophage results in the production of shorter pili, which are associated with both decreased twitching motility and increased phage resistance. This resistance is mediated not only against competing phages, but also phages that are spontaneously produced by the JBD26 lysogen. In fact, the anti-Kronos effect mediated by Zip confers a strong advantage to JBD26 lysogens by allowing large numbers of spontaneously produced phage progeny to accumulate in the culture. These phages can then go on to infect other non-lysogenic cells in the community and thereby promote the spread of phage JBD26. We show that this anti-Kronos effect – protecting the lysogen community from loss of phage progeny by reinfection – is mediated by different prophages using different biological mechanisms. These anti-Kronos proteins provide a strong competitive advantage to prophages that encode them by enabling them to efficiently transmit both vertically as lysogens and horizontally via phage infection.

While the precise mechanism through which Zip inhibits phage infection is not yet known, our data allow us to propose a model. A previous study using cryo-electron tomography suggested that only one type of motor, PilB extension or PilT retraction, can bind to the pilus machinery at any given time^39^. Further work using fluorescence microscopy developed a model where competitive binding of the PilB extension and PilT retraction motors coordinates the repetitive cycles of pilus extension and retraction^40^. This binding process was proposed to be stochastic, with PilB binding to the inner membrane complex and mediating pilus extension before dissociating and opening the binding site to the PilT retraction motor (Figure 1D). Our protein interaction data showed that Zip forms a tripartite complex with PilB/PilZ, and fluorescence microscopy showed that Zip-GFP localizes to the bacterial cell pole, where the pilus is known to assemble. These data suggest that PilB/PilZ/Zip together bind to the pilus assembly site in the inner membrane (Figure 3E). We believe that the addition of Zip to this complex destabilizes the interaction of PilB with the pilus assembly complex, causing it to spontaneously dissociate more rapidly than in the absence of Zip. This activity would result in the formation of shorter pili, consistent with what we observed in the JBD26 lysogen (Figure 1H).

Further support for this model is provided by the dynamics data collected on the JBD26 lysogen, which showed that while the pilus extension velocity was the same as wild-type cells, the pilus extension time was decreased (Figures 1I,J), consistent with the PilB motor prematurely dissociating. Why shorter pili might increase phage resistance is not entirely clear, and this work may hint at a role for the pilus inner membrane complex and/or PilB assembly ATPase in the phage genome entry process. In contrast to *E. coli* phages, for which several inner membrane proteins are known to be important for phage genome entry^16,50–52^, there are no confirmed inner membrane protein requirements for *Pseudomonas* phages. As PilZ is highly conserved across all *P. aeruginosa* strains, it provides an excellent target to modulate to provide resistance to a broad spectrum of pilus-dependent phages.

Zip expression is controlled by LasR, the principal regulator of quorum sensing in *P. aeruginosa*, allowing it to be expressed in a cell density-dependent manner. This permits the lysogen to tune expression in concert with the relative risk of phage infection. At low cell density, there is little risk of being reinfected by phage progeny produced by spontaneous induction from individual bacteria as the cells are spread out. As cell density increases, the chance of infection also increases as the absolute number of cells spontaneously producing phages increases, and the cells within the community are closer together. Thus, it is highly beneficial for the lysogens to respond to this increased risk by modulating expression of their pili and thereby decreasing the chance of phage progeny reinfecting the lysogen. This defence is also protective against other phages that require the type IV pilus for infection that might invade the bacterial community and initiate an epidemic. This quorum-mediated control of anti-phage defence is similar to what has previously been observed for bacterially encoded anti-phage defences. For example, in *P. aeruginosa,* LasR contributes to high cell density-mediated expression of anti-phage defence systems both directly by upregulating the production of a community-based anti-phage defence molecule known as PQS^53^ and indirectly by upregulating expression of RhlR, which drives expression of CRISPR-Cas systems^54^. While phages have been shown to integrate quorum sensing into their lysis-lysogeny decision^56^, to our knowledge, this work provides the first evidence that prophages have connected into this conserved bacterial circuitry to control expression of their antiviral defence genes. As quorum sensing systems are broadly distributed across bacteria, this is likely a widespread feature of prophage-encoded defences.

Previous studies investigating superinfection exclusion and its effects on viral populations showed that allowing superinfection yields populations that are more capable of adapting to changes in the environment as a result of the potential for exchange of genetic material between viruses^57,58^. However, mutants that can prevent superinfection have also been shown to display a significant advantage over their superinfection permissive counterparts, even when this ability came with a substantial cost to their growth rate^57^. Together, these results suggest that while superinfection exclusion is a winning strategy in the short term, preventing superinfection can negatively impact the long-term prospects of a viral population. The quorum mediated control of a superinfection exclusion protein like Zip allows the modulation of activity in a manner that provides anti-phage defence in some environments and permits uptake of genetic material in others, thereby granting both evolutionary benefits. For example, in spatially structured environments, the competition for hosts is local, and lysogens are more likely to be battling genetically identical phages released by nearby cells. In this case there is no advantage to allowing phages to enter the lysogen. Zip expression would be upregulated in direct measure with cell density in the structured environment and exclude these phages from infection, thereby also enabling higher numbers of free phages to escape the structured community in search of new hosts. By contrast, in well-mixed environments where competition is global and lysogens are more likely to encounter genetically diverse phages, low Zip expression could allow other pilus-dependent phages to infect, thereby providing the advantage of genetic exchange. Thus, the measured control of Zip activity provides a means by which prophages can balance these two opposing evolutionary advantages.

The success of a virus depends on its ability to spread both within and between hosts^59^. With bacterial viruses, the spread within host is provided via vertical transmission of the prophage from mother to daughter cells that naturally occurs when cells divide. Spread between hosts involves the horizontal movement of the phage to uninfected cells. For this to efficiently proceed, the phages produced by a lysogenic community need to avoid cells that already contain a prophage and efficiently seek out cells in which they can initiate a fruitful infection. Zip activity promotes this by inhibiting the ability of phages to bind to lysogens and reinfect. We call this the anti-Kronos effect. We have shown that this effect has a large impact on free phage numbers produced by lysogens for two other temperate phages that are unrelated to JBD26 and use different mechanism to block cell surface binding by phages. We expect that the anti-Kronos effect will manifest in many other diverse temperate phage systems. This property of viral self-exclusion has also been described in eukaryotic viruses like HIV, where Nef downregulates CD4+, the viral receptor to support viral propagation, and vaccinia virus, where viral proteins were shown to mediate repulsion of virions from the surface of infected cells, thereby speeding up viral spread^9,60,61^. Our work shows that this common eukaryotic virus trait is conserved among bacterial viruses, providing another example where the interaction of bacterial viruses and their hosts is mirrored in eukaryotic systems^62–64^.

## Data and code availability

This paper does not report original code.

Any information required to reanalyze the data reported in this paper is available from the lead contact upon request.

## Acknowledgments

The authors thank members of the Maxwell, Davidson and Gitai laboratories for helpful discussions. This study was supported by grants from the Canadian Institutes of Health Research to K.L.M. (PJT-165936) and A.R.D. (FDN-15427), and a Natural Sciences and Engineering Research Council Arthur B. McDonald Fellowship to K.L.M. (SMFSU-581368-2023). A.R.D. is the Canada Research Chair in Bacteriophage-Based Technologies. V.L.T. is supported by a Career Transition Award granted by the Emerging Pandemic & Infections Consortium (EPIC) at the University of Toronto.

## Author contributions

V.L.T. and K.L.M. conceptualized the project. Phage experiments, genome deletions, plasmid construction, protein purification, bacterial-two hybrid assays, and motility assays were performed by V.L.T., with assistance from M.S. and P.H.P.. C.M.B. performed the fluorescence localization assays. A.Y. and M.D.K performed the i*n vivo* pilus dynamics and quantification. Z.G., A.R.D., V.L.T., M.D.K. and K.L.M contributed to experimental design. The manuscript was written by V.L.T. and K.L.M., and all authors contributed to editing the manuscript and support the conclusions.

## Star Methods

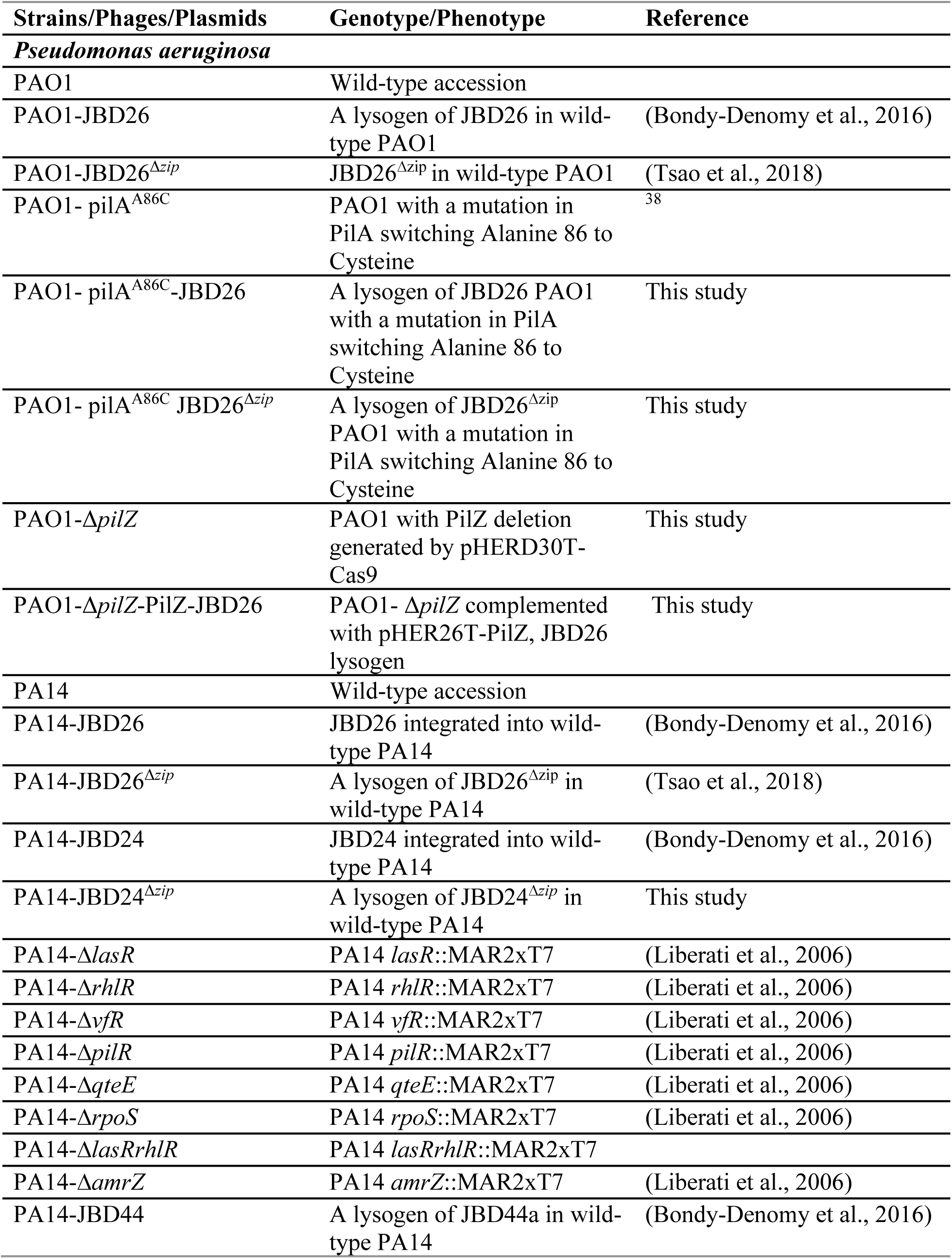
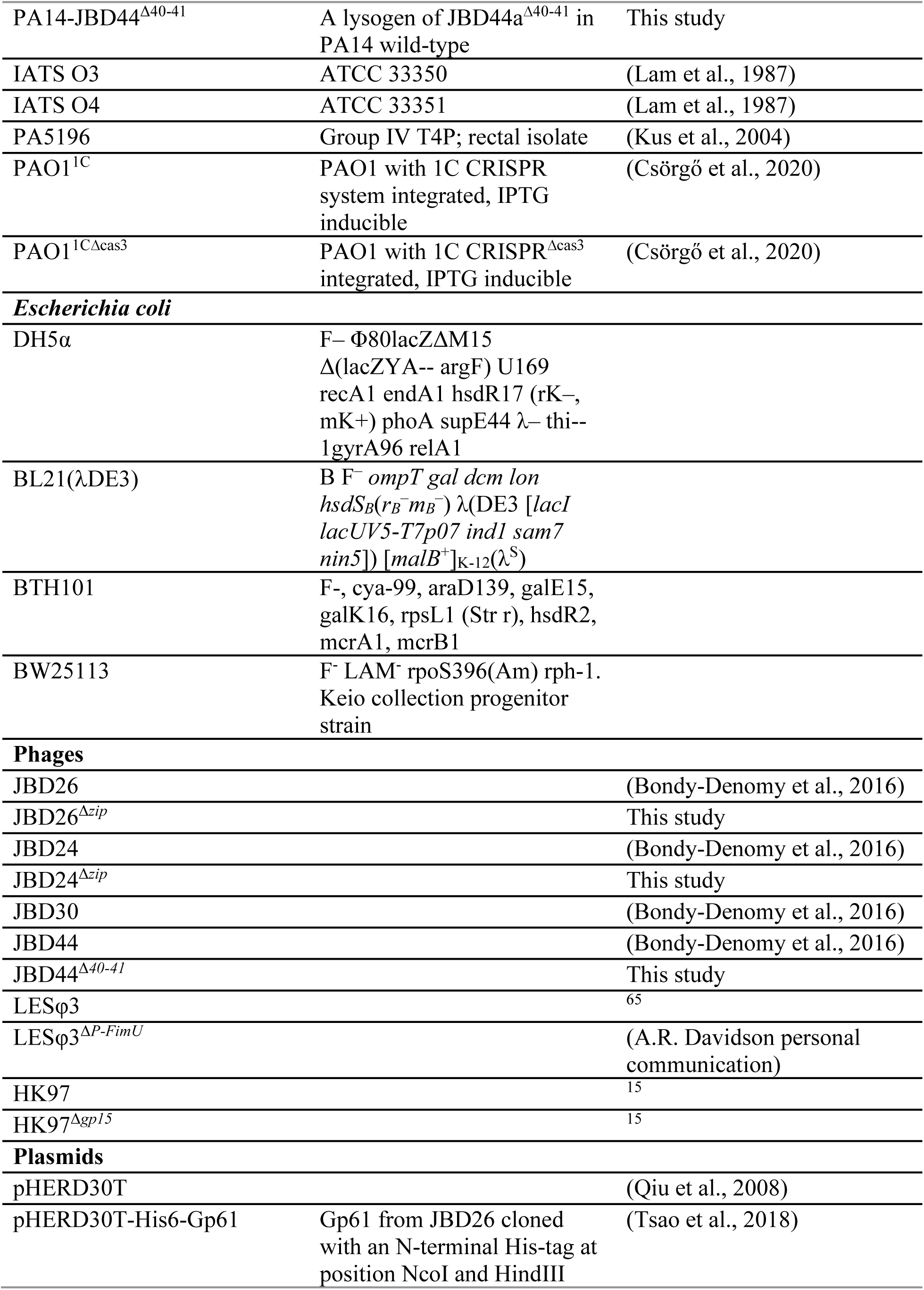
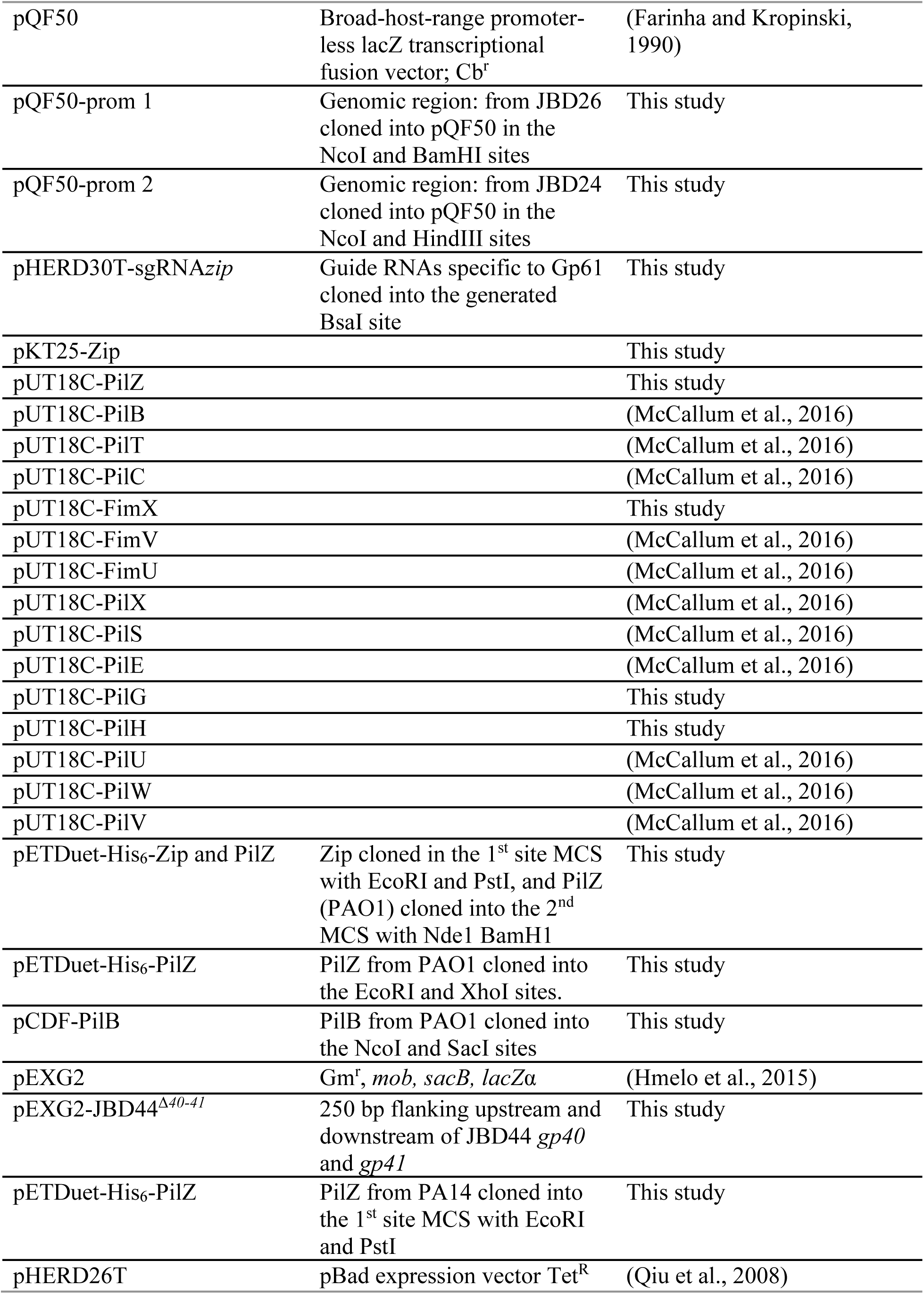
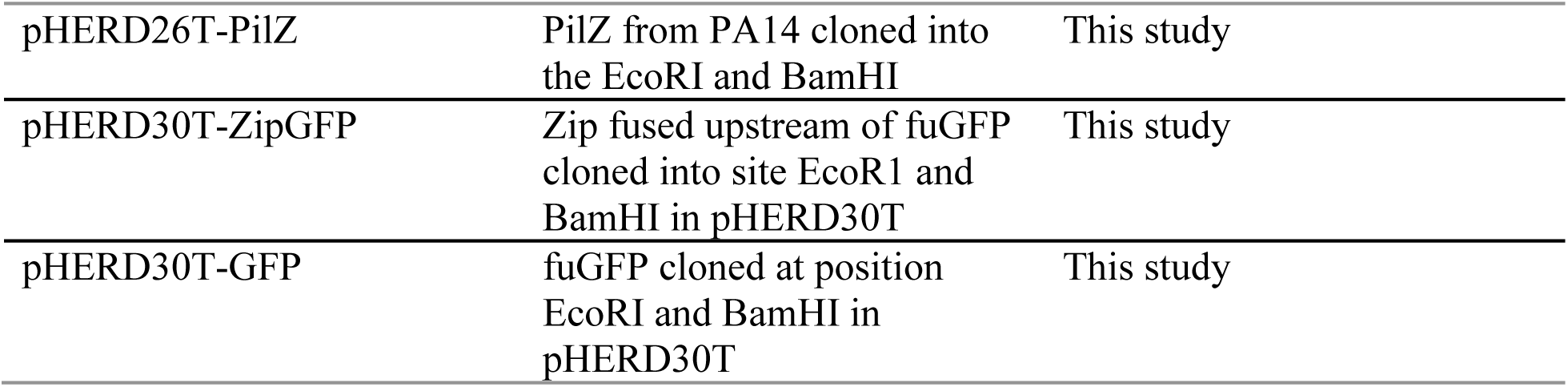

### Resource availability

#### Lead contact

Further information and requests for resources and reagents should be directed to and will be fulfilled by the lead contact, Karen Maxwell.

### Experimental model and subject details

#### Bacterial strains and phages

*E. coli* NEB 5-alpha (New England Biolabs) was the cloning strain used. *E. coli* BL21(λDE3) was used for protein expression and purifications, *E. coli* BTH101 was used for bacterial two-hybrid assays and *E. coli* BW25113 was used for phage-related experiments. *Pseudomonas aeruginosa* strains were obtained from a previous study ^22^. Unless otherwise stated the cells were grown in Lysogeny Broth (LB) with or without 10 mM MgSO_4_, with or without 0.1% L-arabinose and 1.2 or 1.5% agar. Specific antibiotic concentrations used for *E. coli* were: ampicillin (100 μg/mL), tetracyline (12 μg/mL), gentamicin (20 μg/mL) and streptomycin (34 μg/mL) and for *P. aeruginosa*: carbenicillin (300 μg/mL), gentamicin (50 μg/mL) and tetracycline (90 μg/mL). Strains generated for this study and phages used are detailed in the Key Resources table.

#### Identification of tfpZ within the IATS O3 genome

A member of the type V pilus family was identified by analyzing the genomic region between *nadC* and *pilB* of the 20 IATS strains using RAST SEEDViewer server (Aziz et al., 2008), for the presence of *pilA* and *tfpZ*. Polymerase chain reaction (PCR) using primers specific to *tfpZ* were used to confirm presence of the gene.

#### Twitching assay

Individual colonies grown on LB agar supplemented with 50 μg/ml gentamicin and 0.1% L-arabinose were stab inoculated into a 1% LB agar plate supplemented with 50 μg/ml gentamicin and 0.1% arabinose and were incubated for 24 h at 37°C. Twitching motility was assessed by removing the agar and staining the cell mass adhered to the petri plate using 1% crystal violet. Twitching assays with wild-type *P. aeruginosa* and the lysogens were performed in the absence of antibiotics.

#### Phage preparations

High titer phage lysates were obtained by growing *P. aeruginosa* lysogens overnight in LB medium at 37°C. The cells were then diluted 1/100 into fresh LB medium and grown with shaking at 37°C to an OD_600_ of 0.5;mitomycin C was then added to a final concentration of 3 μg/ml, and the cultures were incubated until they lysed. Cellular debris was removed by centrifugation at 15,000 x g for 10 min. The supernatant containing the phages was transferred into a fresh test tube and a few drops of chloroform were added for storage at 4°C. Phage titers were determined by plating ten-fold serial dilutions on plates containing 150 μL bacteria resuspended in 3 ml of 0.7 % top agar supplemented with 10 mM MgSO_4_ (plus relevant antibiotic and 0.1 % L-arabinose when noted). Plates were incubated overnight at 30°C to allow plaques to form.

#### Generating Zip mutants in JBD26, JBD24 and JBD44

To create Zip knockouts in JBD26 and JBD24 a conserved sequence within *zip* was chosen to generate a guide RNA specific to the 1C CRISPR-CAS system. The guide RNA was cloned into pHERD30T and transformed into the PAO1^1C^ and PAO1^1CΔcas3^ strains (Csörgő et al., 2020). To generate the mutations, lysates of JBD26 and JBD24 were isolated from lysogens within PAO1 grown overnight. PAO1^1CΔcas3^ expressing the guide RNA targeting *zip* was grown at 37°C overnight in the presence of 50 µg/ml of gentamicin, 0.1 % L-arabinose and 0.5 mM IPTG. The following morning the culture was diluted 1/100 in the same media and was grown at 37°C until mid-exponential stage. Ten microliters of JBD26 or JBD24 were added to the culture, and it was incubated overnight at 37°C. The following morning supernatants containing the replicated phages were plated on lawns of PAO1^1C^ expressing the guide RNA targeting *zip* with the LB plates and top agar supplemented with the same selection and inducing agents. Following overnight incubation, individual plaques were resuspended in LB and three plaque purification steps were performed. Mutant phages were confirmed by sequencing.

To delete the proposed serotype converting genes *40* and *41* from phage JBD44, a deletion construct was generated to remove the region between nucleotides 28672-31193. We created a construct in pEXG2 (Hmelo et al., 2015) that had 250 base pairs flanking either end of this region to use for homologous recombination. This construct encoded the first 15 amino acids of Gp40 and the final 15 amino acids of Gp41. This pEXG2 construct was transformed into *E. coli* SM10 and mated into a JBD44-PA14 lysogen for homologous recombination. Mutants were confirmed by DNA sequencing.

#### Phage production over time

Three independent lysogens were grown overnight at 30°C with shaking. The following morning bacteria from 100 μl aliquots of the cell cultures were collected by centrifugation and washed three times with 1 mL of LB to remove all free phages. Equal numbers of cells were resuspended in a final volume of 2 mL LB supplemented with 10 mM MgSO_4_ and were incubated at 30°C with shaking.

Samples were taken at 3, 6, and 20 hours; the bacterial cells were collected by centrifugation and serial dilutions were plated onto LB agar and incubated at 37°C overnight to enumerate the number of bacteria present at each time point, and the titers of the phages present in the supernatant fraction were determined through plating assays on PA14.

#### Zip promoter assay

The intergenic region between gene *60* and *zip* was amplified from phage JBD26 and cloned into the NcoI/HindIII sites of the promoterless ꞵ-galactosidase reporter shuttle vector pQF50. To analyze promoter type 2, the intergenic region between genes *57* and *58* was amplified from phage JBD24 and cloned into the NcoI/HindIII sites of pQF50. The constructs were transformed into the designated strains and assayed for β-galactosidase activity. Individual colonies were used to inoculate overnight cultures of LB containing 300 µg/ml of carbenicillin. The following morning cells were diluted into fresh LB with carbenicillin and grown at 37°C to an OD of 0.5 for exponential phase experiments, or 16 h to reach saturation (OD_600_ was measured at this point). β-Galactosidase activity was quantified by mixing 100 μL of culture with 900 μL of Z buffer (0.06 M Na_2_HPO_4_, 0.04 M NaH_2_PO_4_-H_2_O, 0.01 M KCl, 0.001 M MgSO_4_ and 0.05 M β-mercaptoethanol), the cells were lysed with 100 μl of 1% SDS and 100 μl chloroform and vortexed for 10 sec. The cultures were incubated at 30°C for 5 min and 200 μl of 40 mg/ml o-nitrophenyl-β-galactosidase resuspended in Z buffer was added before incubating for 15-30 min. The reaction was stopped with the addition of 500 μl 1M Na_2_CO_3_, A_420_ and A_550_ were measured, and the Miller Units were calculated.

#### Zip cellular localization

To determine the cellular localization of Zip at native expression levels, a fusion construct of the *zip* promoter sequence and free-use green fluorescent protein (fuGFP) fused to the N-terminus of *zip* was cloned into pHERD30T. Twitching inhibition and phage resistance of cells expressing prom-fuGFP-Zip were measured. In tandem fuGFP alone was cloned into the pHERD30T construct. The cultures were inoculated on plates containing 50 µg/ml of gentamicin and 0.1% L-arabinose and were grown overnight at 37°C. The following day a single colony of each strain was resuspended in 10 µl of 1X PBS and 2 µl was spotted on 2% agarose pads. For all microscopy, Zip localization was assessed using differential interference contrast (DIC) microscopy or using the enhanced green fluorescent protein (EGFP) channel on a Zeiss Axio Imager M1 microscope (Carl Zeiss) at the same exposure time. Three independent fields were captured for each sample and images are representative of three technical replicates across two biological replicates.

#### Pilus dynamics experiments

Pilus labeling, data acquisition, and data analysis were performed as described previously (Koch et al., 2022). Briefly, cells were grown overnight and used directly in high cell density experiments or diluted 1:1000 and grown to mid-log phase (OD = 0.3) for low cell density experiments. Cells were incubated with 35 ng/µl Alexa Fluor 488 maleimide dye for 45 minutes to stain pili. Unbound dye was washed away by pelleting cells twice and resuspending in fresh medium. Cells were spread on an agarose pad covered with a coverslip for imaging under a Nikon TiE or Nikon Ti2 fluorescent microscope with a 100x NA1.45 Ph3 objective lens. Videos of pilus dynamics were recorded for 30 s each. Pilus dynamics were analyzed in ImageJ ^66^.

#### Bacterial two-hybrid assay

To identify an interaction partner for Zip, we used a library of pilus biosynthesis genes cloned into the Euromedex bacterial two-hybrid system (Battesti and Bouveret, 2012). The *E. coli* BTH101 strains lacking a functional adenylate cyclase were transformed by pKT25-zip and individual pilus genes cloned into pUT18C. The resulting transformants were plated on LB agar plates containing ampicillin (100 µg/mg) and kanamycin (50 µg/ml). The plates were incubated at 30°C overnight and three individual colonies were selected per plate to assay for interactions. LB cultures were grown overnight at 30°C in the presence of the stated ampicillin and kanamycin concentrations and supplemented with 0.5 mM IPTG. The next morning 2 µl of culture was spotted onto two sets of chromogenic plates: LB agar with X-gal (40 µg/ml) and MacConkey agar, both of which contained ampicillin (100 µg/ml), kanamycin (50 µg/ml) and 0.5 mM IPTG. The plates were incubated for 24 h at 30°C. The presence of a blue colony (X-gal) or a red colony (acidification of MacConkey medium) indicates a positive interaction. The overnight culture was then diluted 1/100 in fresh supplemented media and grown at 30°C with shaking for 6 h. The Miller units were obtained by assaying for β-galactosidase activity as described above.

#### Protein expression and purification

Single expression of His_6_-Zip was achieved used p15TV-L (Genbank: EF456736) and PilZ was cloned in site 1 of pETDuet in frame with the N-terminal hexa-histidine (6His) tag. For co-expression, genes were cloned into pETDuet with Zip in MCS-1 in-frame with the N-terminal 6-His tag and PilZ in MCS-2. Plasmids were propagated in *E. coli* DH5a supplemented with 100 µg/ml of ampicillin. To express three proteins at once, PilB was cloned into pCDF1-b using the NcoI and SacI restriction sites to remove the 6His-tag. Co-expressions with pETDuet His_6_-Zip-PilZ and pCDF-PilB were supplemented with 100 µg/ml of ampicillin and 34 µg/ml streptomycin. Purifications were performed from 2 L cultures of *E. coli* BL21(λDE3) as outlined below.

The cells were grown at 37°C with shaking until the OD600 reached ∼0.8, at which point the culture was induced with 1 mM IPTG and the temperature decreased to 20°C for expression overnight. The following morning cells were harvested by centrifugation and resuspended in 200 ml of binding buffer: Tris-HCl pH 8.0 with 250 mM NaCl. The cells were lysed by sonication and cellular debris was removed by centrifugation at 20,000 x g for 20 min. The lysate was incubated with Ni-NTA resin and washed with binding buffer containing 30 mM imidazole and eluted with 250 mM imidazole. The elution fractions were separated on 15% Tris-Tricine gels.

