## Supplemental Figure 1 and Supplemental Figure 2 for "Prophages block cell surface receptors to ensure survival of their viral progeny"

### Slide 1
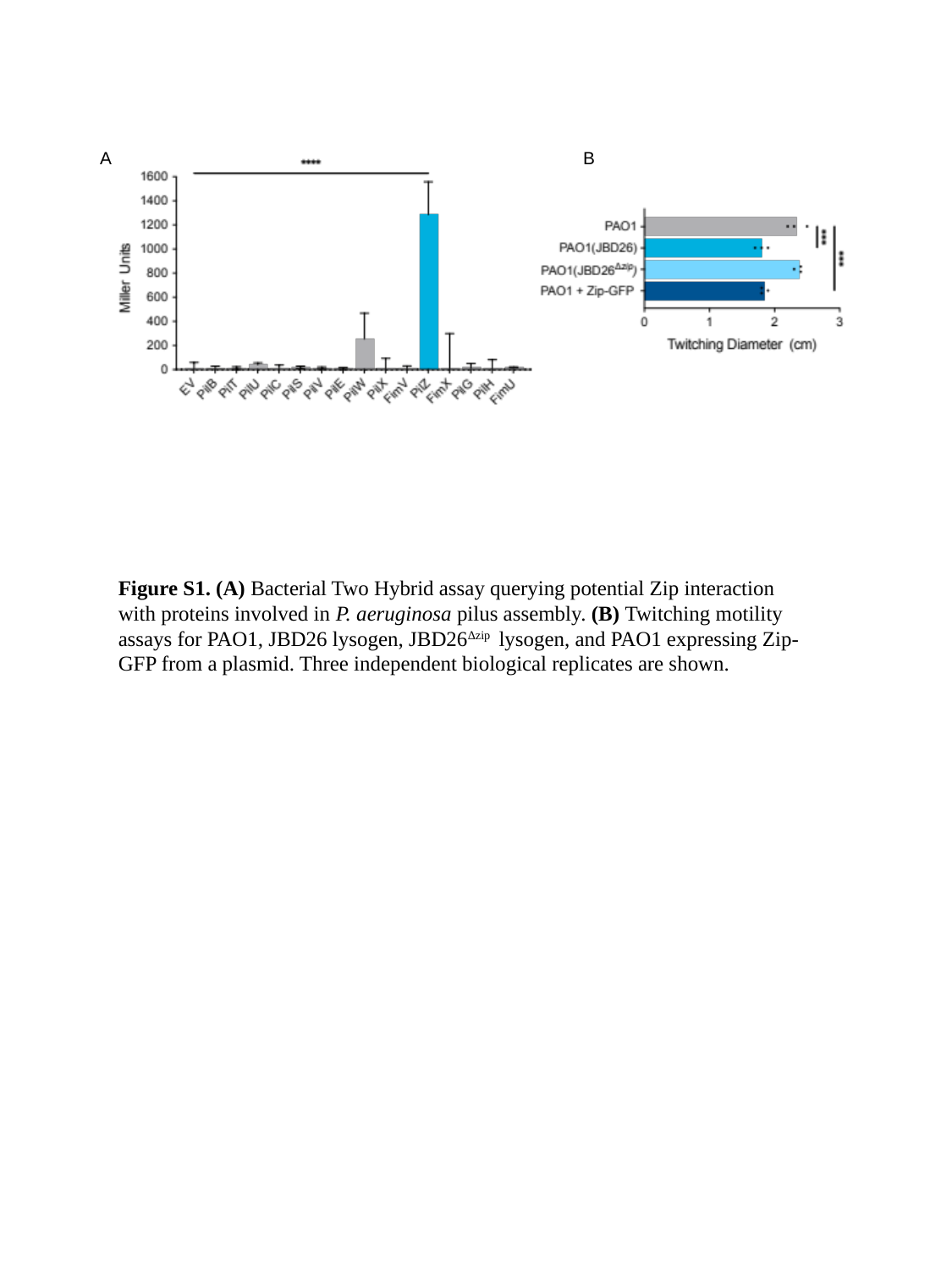

B
A
Figure S1. (A) Bacterial Two Hybrid assay querying potential Zip interaction with proteins involved in P. aeruginosa pilus assembly. (B) Twitching motility assays for PAO1, JBD26 lysogen, JBD26^∆zip^ lysogen, and PAO1 expressing Zip-GFP from a plasmid. Three independent biological replicates are shown.

### Slide 2
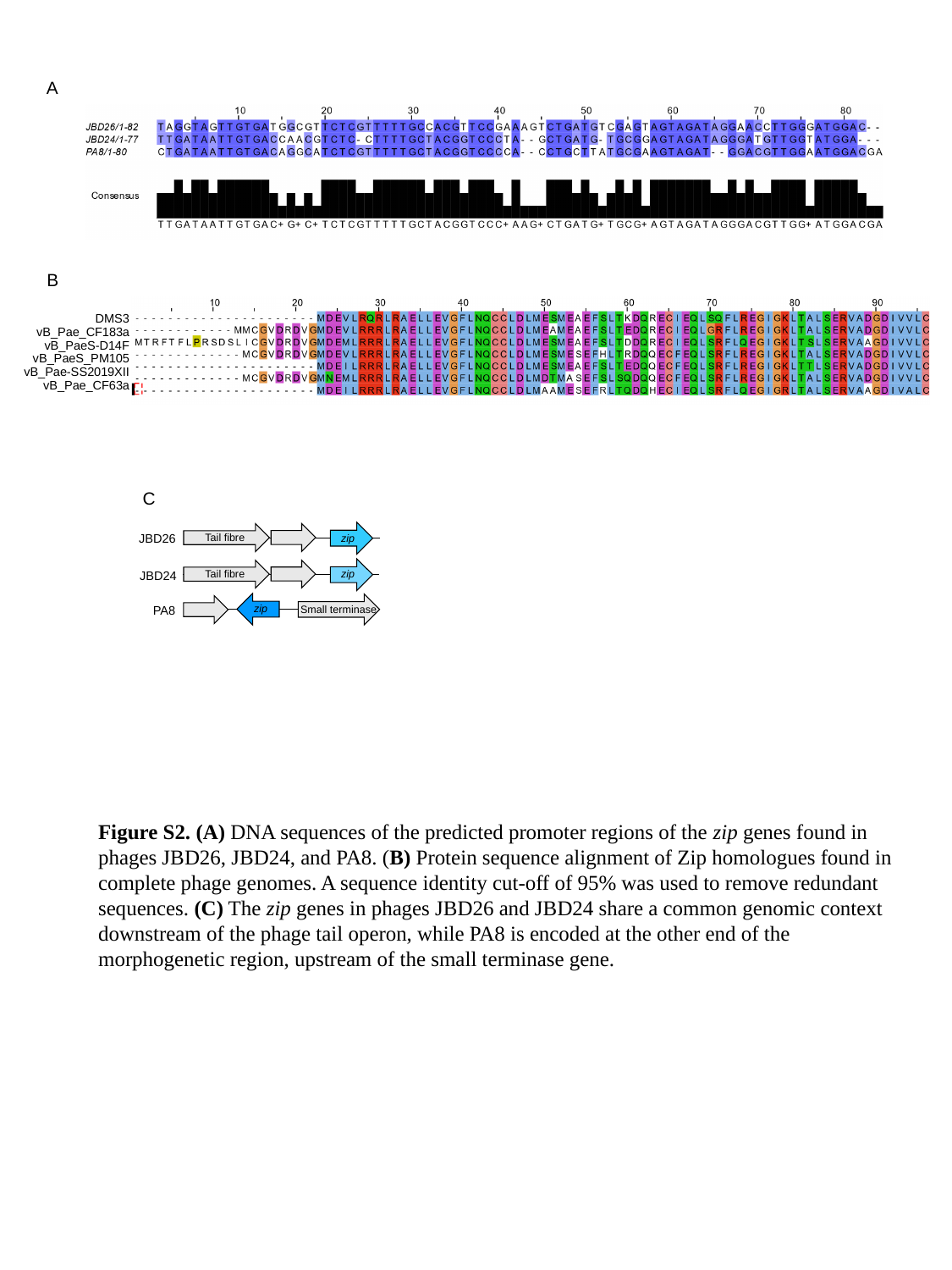

A
B
DMS3
vB_Pae_CF183a
vB_PaeS-D14F
vB_PaeS_PM105
vB_Pae-SS2019XII
vB_Pae_CF63a
C
Tail fibre
zip
Tail fibre
zip
zip
Small terminase
JBD26
JBD24
PA8
Figure S2. (A) DNA sequences of the predicted promoter regions of the zip genes found in phages JBD26, JBD24, and PA8. (B) Protein sequence alignment of Zip homologues found in complete phage genomes. A sequence identity cut-off of 95% was used to remove redundant sequences. (C) The zip genes in phages JBD26 and JBD24 share a common genomic context downstream of the phage tail operon, while PA8 is encoded at the other end of the morphogenetic region, upstream of the small terminase gene.
